# Developmental effects of barium in planaria

**DOI:** 10.1101/2024.02.15.580538

**Authors:** Alberto Molano

**Author notes:** https://www.researchgate.net/profile/Alberto-Molano.

## Abstract

An investigation was carried out of the stage-specific effects of barium, a wide spectrum potassium channel blocker that depolarizes cells, on planaria embryologic development. Barium exposure triggered a variety of axes abnormalities in planaria, including dorsoventral malformations (formation of pigmented dorsal bulges, suggestive of BMP up-regulation), anteroposterior abnormalities (two-headed worms with two eyes or a single eye on each end, worms with an ectopic anterior partial body embedded diagonally within the “main” body of the planaria), and other congenital anomalies (baby worms with extremely short bodies, worms with multiple eyes or with a strange tubular structure emerging from the malformed eyes). The implications of these findings for current understanding of planaria development are discussed.

## Introduction

Recent evidence supports the notion that ion channels play a role during embryogenesis. For example, a gain-of-function mutation in the CACNA1C gene, which codes for the L-type voltage-gated Ca^2+^ channel Ca_v_1.2, causes Timothy syndrome, a rare disorder characterized by cardiac arrythmias, cutaneous syndactyly, facial dysmorphia, and baldness at birth [1]. Although the heart conduction anomalies have a straightforward explanation, the developmental abnormalities affecting non-excitable tissues suggest unexpected roles for ion channels during embryogenesis.

Similarly, mutations in the KCNJ2 and KCNJ5 genes, which code for the inwardly rectifying potassium channel Kir2.1, and the G-protein-activated inwardly rectifying potassium channel Kir3.4, respectively, cause Andersen–Tawil syndrome, a rare autosomal dominant disorder characterized by recurrent flaccid muscle weakness (periodic paralysis), cardiac arrhythmias, and distinctive skeletal and facial features which include low-set ears, broad forehead, ocular hypertelorism, small mandible, dental abnormalities, fifth digit clinodactyly, syndactyly, short stature, and scoliosis [2]. The presence of these cranial, skeletal, facial, dental, and finger anomalies reinforce the notion that ion channels play a role during embryogenesis.

Animal models have begun to provide detailed molecular, stage-specific evidence which supports the human genetic data. For example, it was shown that KCNJ2 is expressed in *Xenopus laevis* during the earliest stages of craniofacial development, and that injecting the embryos with mRNA encoding the same mutations found in Andersen–Tawil human patients caused comparable craniofacial abnormalities [3].

From a pharmacologic standpoint, intrauterine exposure to anti-epileptic drugs that inhibit ion channels is likewise associated with increased incidence of congenital malformations [4]. Examples include topiramate, valproate, ethosuximide, phenobarbital, phenytoin, and carbamazepine. A case in point is phenytoin, a first-generation anti-epileptic drug that inhibits sodium channels. Exposure during pregnancy can cause developmental abnormalities which include growth deficiency, cleft lip and palate, congenital heart defects, abnormal finger and toenails, genitourinary abnormalities, and neurological impairment [4]. Exposure of zebrafish or chick embryos to anti-epileptic drugs is lethal or teratogenic [5] [6].

Barium is a divalent cation with an ionic radius slightly smaller than potassium. Crystallographic data has demonstrated that its small size allows it to fit into the so-called “selectivity filter” of potassium channels [7]. However, its increased positive charge causes it to bind too tightly, preventing the flow of potassium. Therefore, barium acts as a broad-spectrum potassium channel inhibitor. In addition, barium can mimic some of the effects of calcium. Prenatal exposure to barium is associated with neural tube and congenital heart defects in humans [8]. Intragastric administration of barium can induce neural tube defects in pregnant mice [8]. In chick embryos, barium induces specific and reproducible defects, ranging from achondroplasia to bizarre abnormalities and stunting [9]. Exposure of embryos of the marine bivalve *Mytilus californianus* to barium produced abnormal shell calcification and embryo morphology. In this marine species, stage-specific experiments revealed that gastrulae were the most sensitive, while blastula and trochophore larvae were less so [10].

Here, an investigation was carried out of the stage-specific effects of barium, a wide spectrum potassium channel blocker, on the development of planaria. Ion channels help generate a specific voltage state that can be shared through gap junctions to modulate cell-to-cell communication during embryogenesis [11].

Planaria embryologic development is startlingly different from other species, since blastomeres do no remain in direct contact with one another in the earliest stages [12] [13] [14] [15]. Other peculiarities of planaria development include:

1. Eggs are ectolecithal, meaning that the much-needed nutritious yolk is not incorporated within the oocytes. Instead, it is enclosed in separate cells, called yolk cells, produced by a separate organ (the *vitellarium*), and which surround the oocytes. As cleavage starts, the yolk cells that surround the zygotes fuse to form a syncytium. This syncytium is surrounded by intact yolk cells.
2. It is impossible to recognize a stereotypical gastrula stage, since blastomeres do not remain attached to each other, forming a compact morula. Instead, they lose contact with each other and migrate actively to various positions within the syncytial yolk. This has been called “blastomere anarchy”. At an early stage, some of these “anarchic” blastomeres differentiate into a motile, yolk-feeding “cryptic larva”, having an epidermis, embryonic pharynx, and neuron-like cells.
3. Within the egg capsule, the cryptic larvae move freely and feed on the surrounding yolk cells. This has been interpreted as embryonic competition, an evolutionary adaptation that perhaps arose when several developing embryos found themselves trapped inside an egg capsule with limited food resources (yolk cells), and where developing embryos are on average each from a different father [15]. As the cryptic larvae feed, their primitive gut cavity fills up, compressing the surrounding yolk syncytium (containing the remaining “anarchic” blastomeres) into a thin peripheral rim.
4. Within this peripheral rim, blastomeres proliferate and differentiate to form the various tissues of the definitive juvenile worm. In contrast, the tissues of the cryptic larvae are resorbed and disappear.

## Materials & Methods

### Planaria

Planaria were collected from Río Sáchica, a stream in Villa de Leyva, Boyacá, Colombia (5.6365° N, 73.5271° W; altitude 2,149 meters above sea level). Morphologically they appear to be *Girardia sp*. Worms reproduced sexually and were maintained in a tank filled with bottled stream water and oxygenated with an aquarium pump. They were fed raw beef liver once a week.

### Eggs

The moment of egg deposition was determined visually by means of frequent inspections and by the color of the capsule (amber immediately after deposition but turning black after a few hours).

### Barium chloride

Barium chloride dihydrate (99% purity, batch # B401332202, calcium max. 0.05%) was purchased from Loba Chemie (Mumbai, India). Fresh 10 mM stock solutions were prepared in bottled water before each treatment.

### Treatments

The planaria embryos studied in this paper take approximately 20 days to develop from deposition to hatching. To investigate the stage-specific effects of potassium channel blockade during development, barium chloride was added during five intervals: (A) Days 1-4, (B) 5-8, (C) 9-12, (D) 13-16, and (E) 17-20. Freshly prepared solutions of barium chloride dihydrate (10 mM, 5 mM, 4 mM, and 3 mM) were added to the eggs (n=8-15) at the specified time points, followed by extensive washing with water. Images were captured with a smartphone and a Swift 380T microscope (Xiamen, China).

## RESULTS

Planaria embryologic development is usually divided into 8 stages [12] [13] [14]. Some critical morphological and gene expression events have been identified in other planaria species like *Schmidtea polychroa* and were used as the basis for this study (Figure 1).

**Figure 1.**
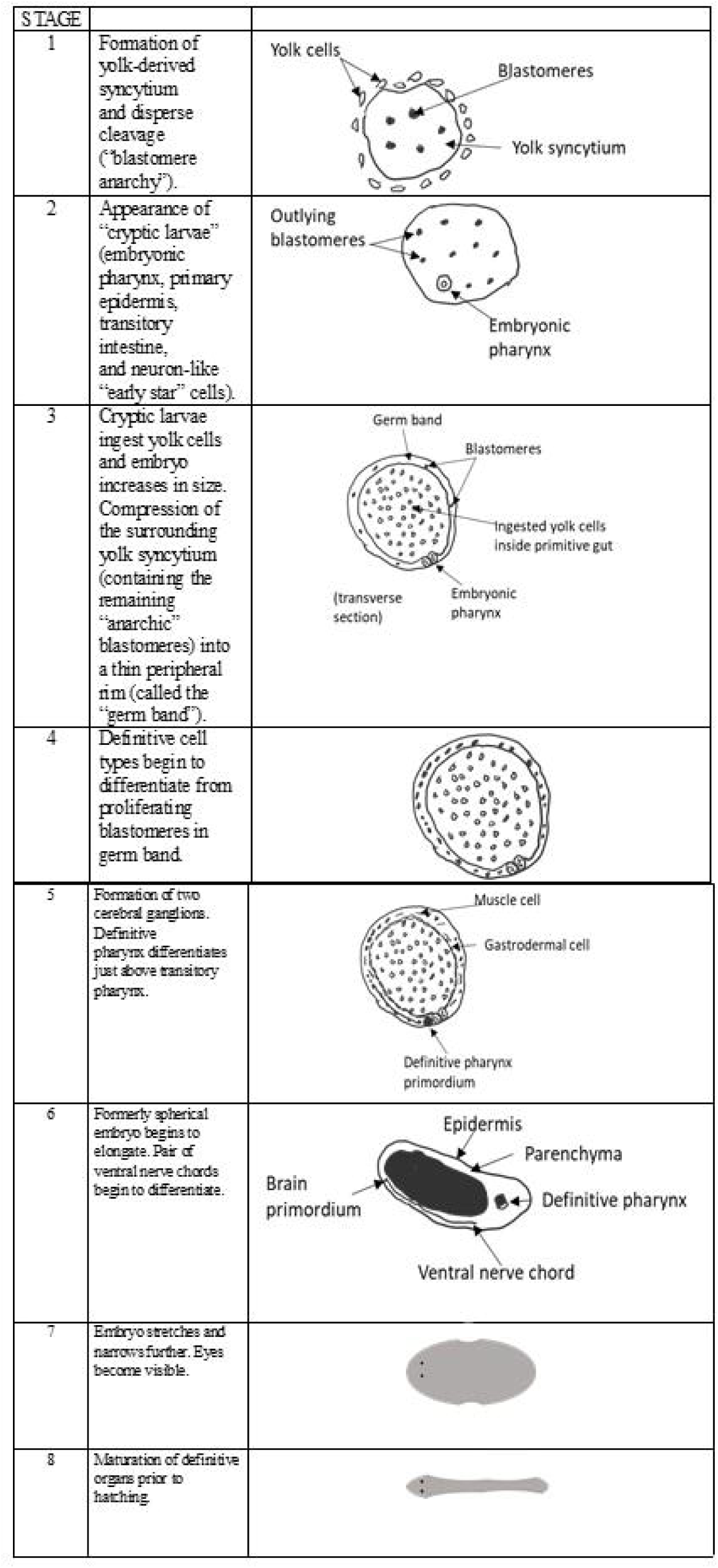
Planaria embryogenesis is usually divided into eight stages. During the earliest stages (1-3), several “cryptic larvae” develop inside the egg capsule through a process known as “blastomere anarchy”, whereby blastomeres do not remain attached to one another but wander freely through the yolk-derived syncytium. These cryptic larvae are eventually resorbed. Definitive cell types begin to differentiate in stage 4 from the proliferating blastomeres located in the peripheral “germ band”. These will give rise to the baby worm.

The planaria embryos studied in this paper hatch 20 days after deposition on average. To match developmental events with days post-deposition, eggs were punctured at various time points (Figure 2). Minutes after deposition, a loose collection of cells could be found inside the capsule, corresponding to lipid-rich vitellocytes. After 4 days, spherical embryos were detected, corresponding to stage 2. After 8 days, embryos of ∼1 mm in size were detected; these were covered with ciliated epithelium that allowed them to slowly glide through the water (video available).

**Figure 2:**
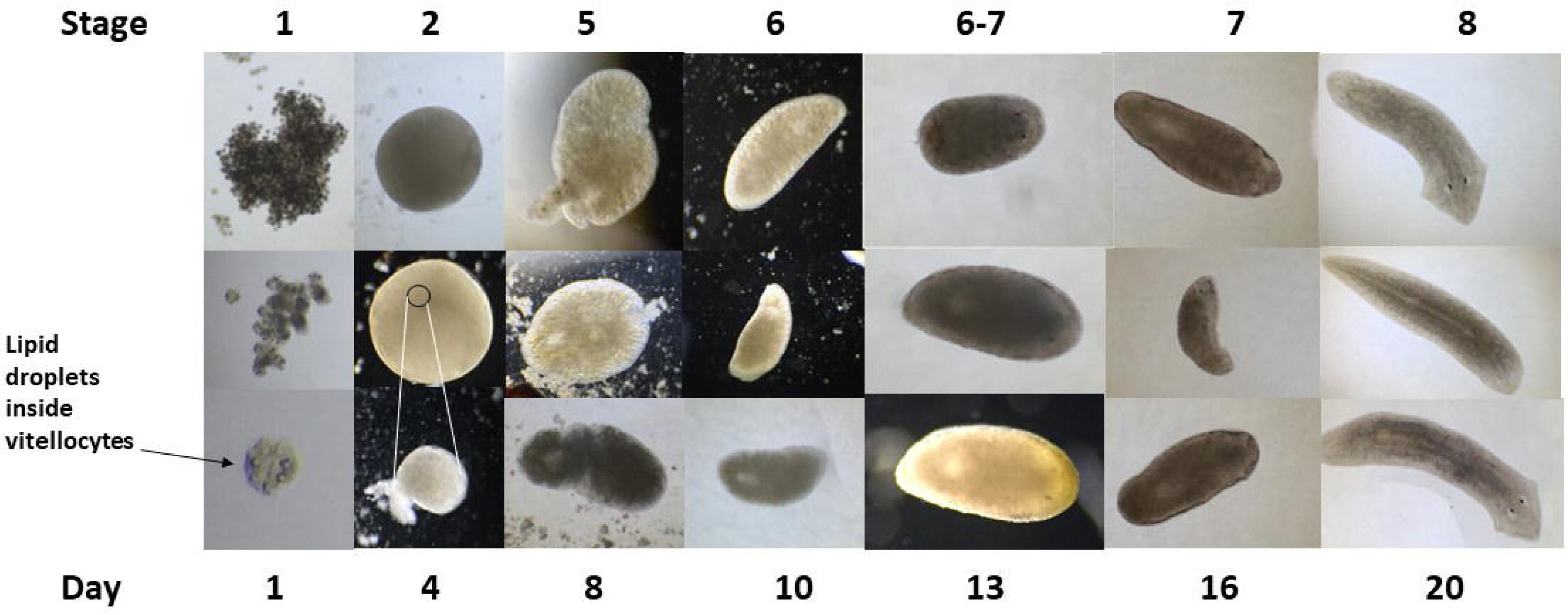
Embryologic stages. Eggs were gently punctured and dissected at the indicated time points/stages to analyze embryo morphology. Notice the lipid droplets inside vitellocytes in stage 1 embryos. The upper two photographs of stage 2 embryos are fixed specimens (4% formaldehyde overnight) showing the entire content inside the egg capsule, whereas the lower picture shows an embryo dissected in an unfixed specimen. The small circle in the middle picture indicates the approximate size of the embryo in relation to the egg.

The Planosphere web tool of the Stowers Institute for Medical Research, which integrates genomic, transcriptomic, phenotypic, and anatomical data of *Schmidtea mediterranea*, was used to investigate potassium channel expression in planaria embryos [16]. Using the search term “potassium channel”, more than 100 transcripts were identified. These transcripts were obtained through RNA-Seq of stage-specific embryos and showed widely different expression patterns during development [17] [18]. Four examples are shown in Figure 3. This analysis indicates that some potassium channels are preferentially expressed during the early (yolk, S2, S3) or late stages (S6-S8), whereas others show both early and late expression, or are expressed throughout development. Please refer to Figure 1 for an overview of planaria embryologic stages (S).

**Figure 3.**
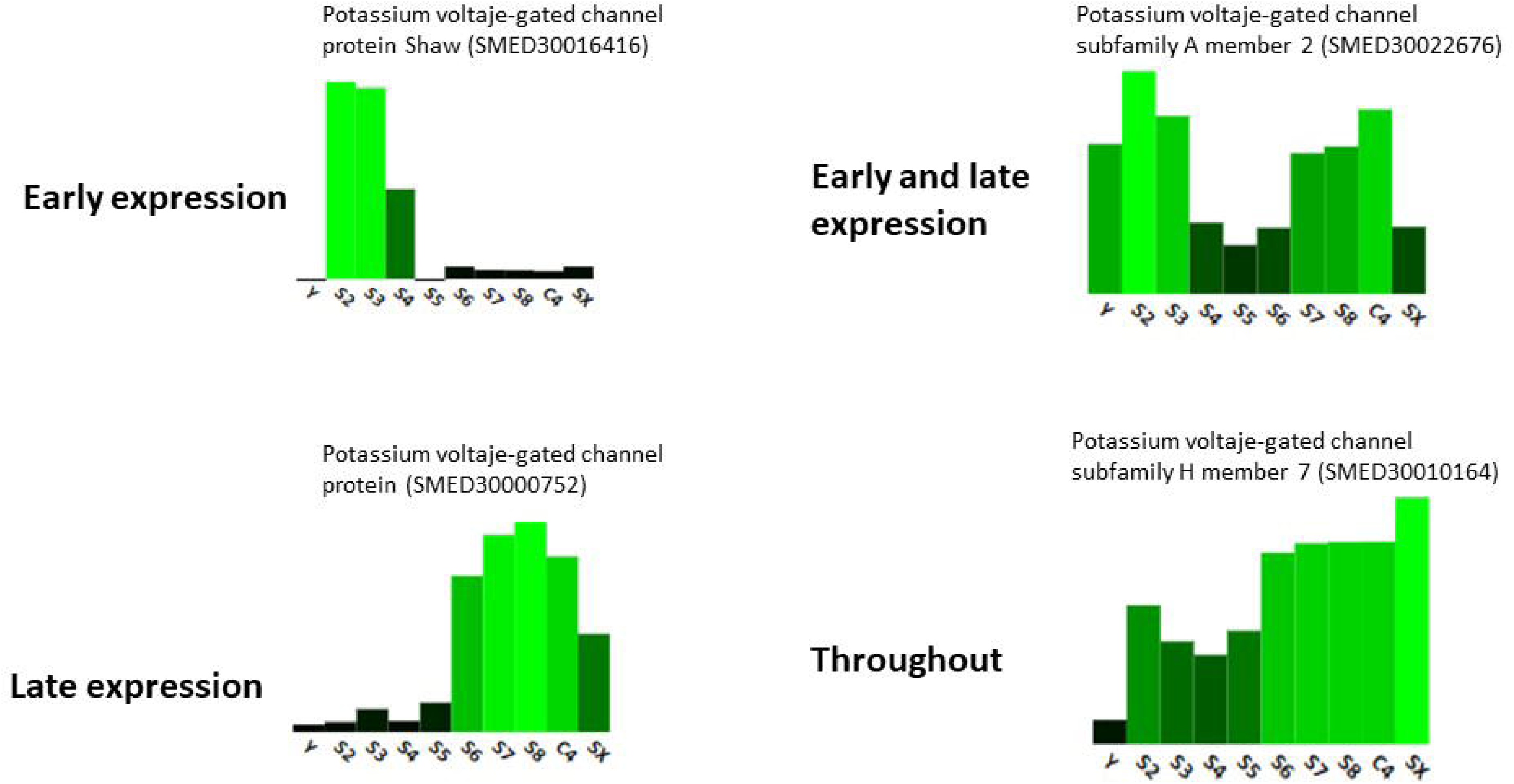
Single embryo RNA-Seq expression analysis of potassium ion channels in the planaria *Schmidtea mediterranea.* Diagram shows four examples of >100 transcripts available through the Planosphere web tool of the Stowers Institute for Medical Research [16] [17]. Bars indicate relative levels of expression during the embryologic and adult stages: Y (yolk), S2-S8 (stages 2 through 8), C4 (asexual adult), SX (virgin, sexually mature adult). Legends above bar graphs indicate the name of the transcript and the Smed ID. Yolk (Y) replicates were prepared from egg capsules lacking developing embryos at 8 days post capsule deposition [18].

To investigate the stage-specific effects of potassium channel blockade during development, barium chloride was added during five intervals: (A) Days 1-4, (B) 5-8, (C) 9-12, (D) 13-16, and (E) 17-20 (see Figure 4). Following barium exposure, eggs were thoroughly washed with water and allowed to continue developing. The following barium concentrations were tested: 10, 5, 4, and 3 mM.

**Figure 4.**
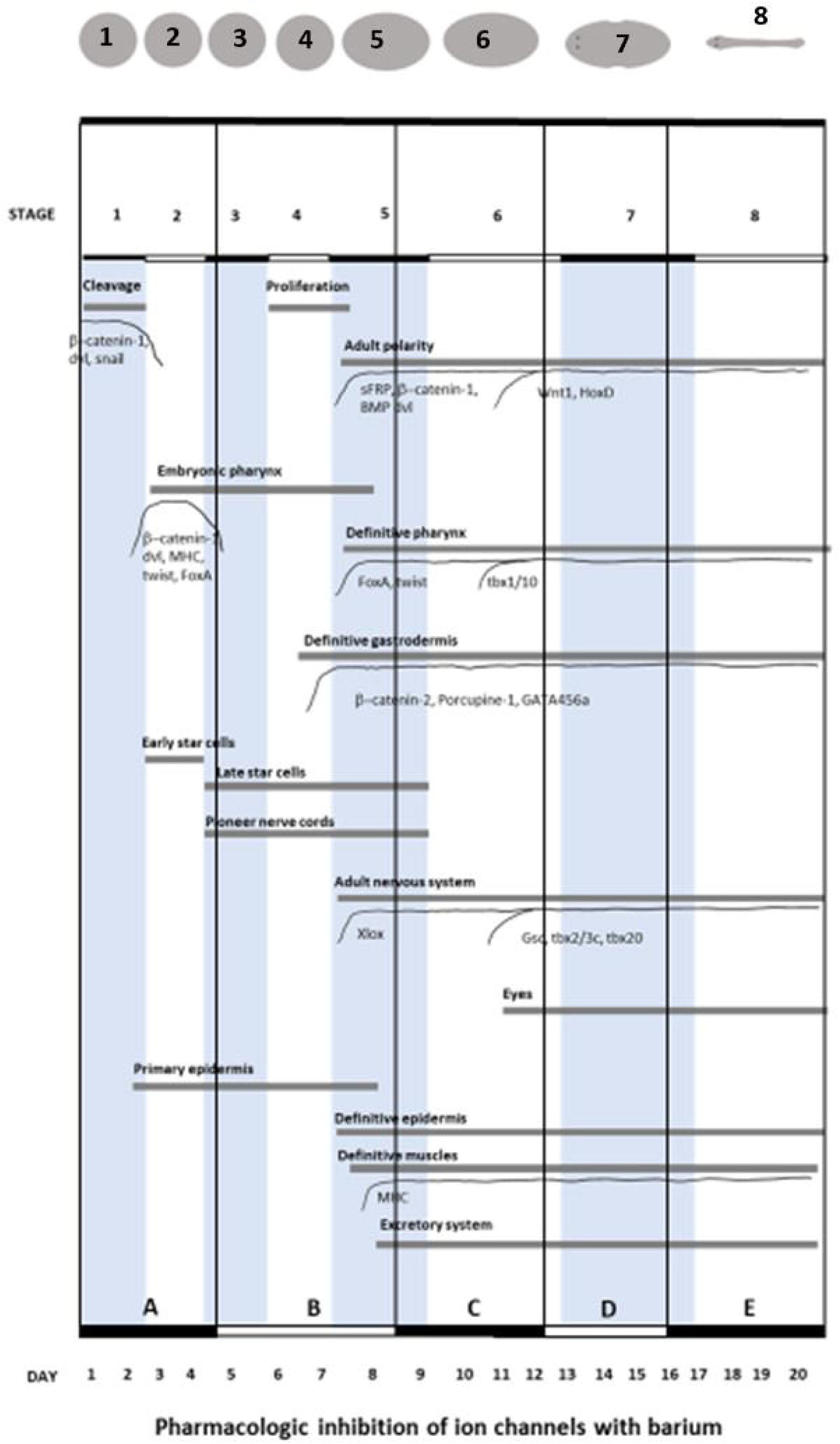
Experimental design superimposed on a diagram of planarian embryonic development [12]. Stages (marked as alternating white and gray stripes) and the approximate shapes of embryos are shown on top. Days are shown on the bottom. Major developmental events are indicated by horizontal bars and associated gene expression by curves. Blastomere cleavage and cryptic larva formation occur in stages 1-3 (∼days 1-6). Definitive cell types begin to differentiate during stage 4 (from about day 6 onwards). Barium was added during the five intervals shown in the lower part of the figure: (A) Days 1-4, (B) 5-8, (C) 9-12, (D) 13-16, and (E) 17-20.

At concentrations ≥ 5 mM, barium was lethal for the earliest stages (figure 5). After puncturing the eggs on day 21 post-deposition, some differences were noted between the two highest barium concentrations tested (10 and 5 mM, data not shown). The highest concentration (10 mM) was lethal and seemed to stall development at a much earlier stage, since the extruded material was like that obtained of an untreated egg punctured immediately after deposition, i.e., a loose collection of cells. This extruded material consisted of dead vitellocytes, without embryo formation (stage 1). In contrast, treatment with 5 mM barium stalled development at a later stage (S2 or S3) since spherical embryos (∼200 – 300 μm in diameter) were clearly visible (data not shown).

**Figure 5.**
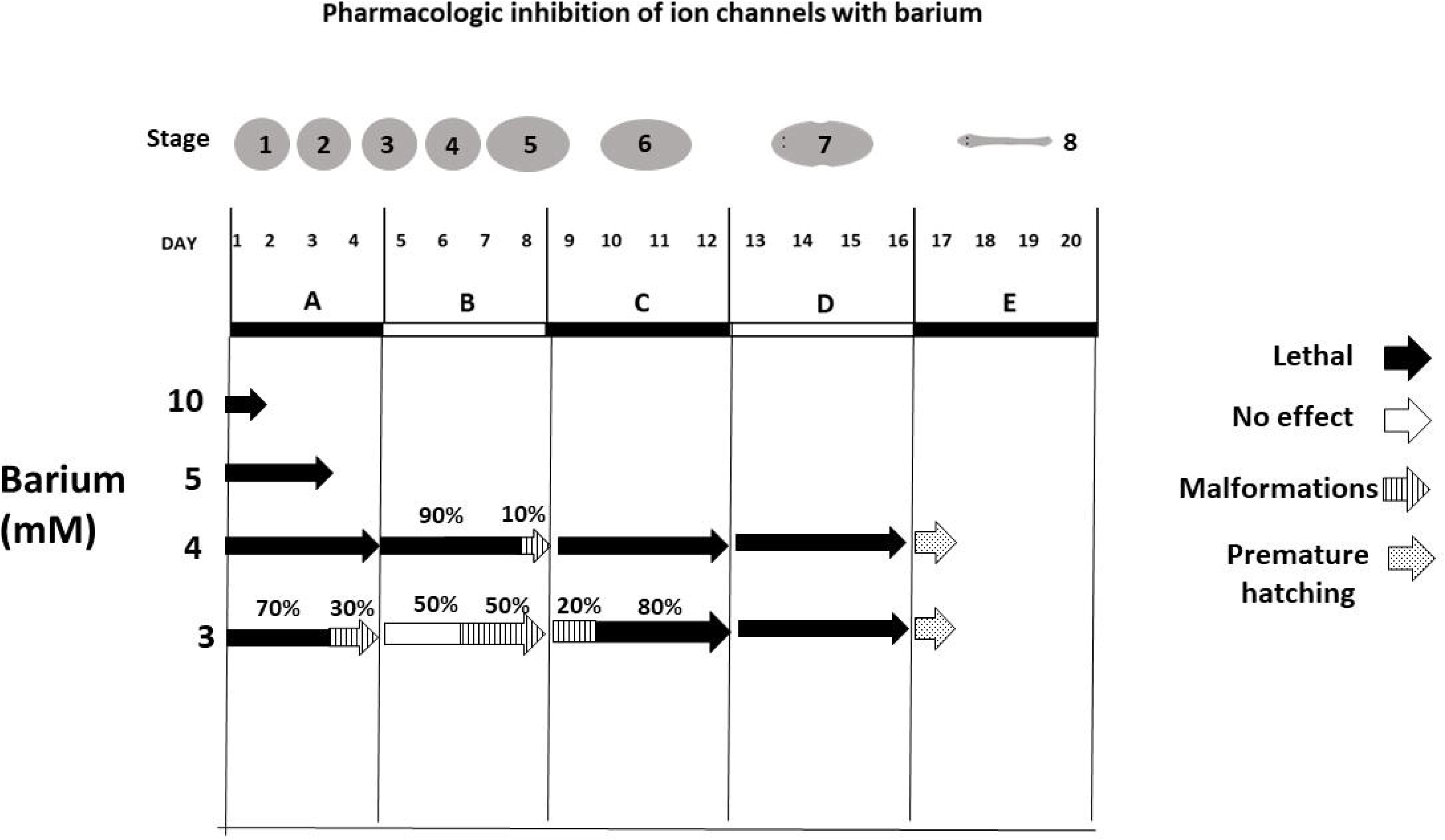
Summary of experimental results. At barium concentrations of 10 mM and 5 mM development stalled in stages 1 and ∼ 2-3, respectively. At 4 mM, barium was lethal after exposure on days 1-4, 9-12, and 13-16. On days 5-8 it very occasionally produced malformations (multiple eyes, not shown), while exposure on day 17 led to premature hatching (within 30 minutes of adding the barium), twitching and disintegration. At 3 mM, barium was lethal or teratogenic on days 1-4 and 9-12 and had no effect or was teratogenic on days 5-8. On days 1-4 barium was lethal for 70% of the eggs and induced malformations in 30%. On days 5-8 barium had no effect in 50% of the eggs and induced malformations in 50%. On days 9-12 barium was lethal for 80% of the eggs and induced malformations in only 20%. On days 13-16 barium was 100% lethal. When 3 mM barium was added to day 17 embryos, premature hatching was triggered within 60 minutes, and it induced twitching, peripheral disintegration, and death.

At 4 mM barium was 100% lethal on days 1-4, 9-12, and 13-16 (Figure 5). On days 5-8 it very occasionally produced malformations, such as multiple eyes. Interestingly, when 4 mM barium was added to day 17 embryos, premature hatching was induced within 30 minutes and barium had a lethal effect on the embryos. Some of the embryos disintegrated [19].

Malformations were seen mainly after exposure to 3 mM barium, and were classified as follows (figure 6):

1. Dorsoventral axis malformations: pigmented dorsal “bulges”. These were seen only after exposure on days 5-8 and 9-12 (but never on days 1-4).
2. Anteroposterior axis malformations. These were seen only after exposure on days 1-4 and 5-8, and included:

a. Two-headed worms (with 2 eyes or 1 eye on each end).
b. Worms with second partial bodies embedded within the “main” body of the planaria, mostly heads.
3. Other malformations: baby worms with multiple eyes, with a strange tubular structure emerging from a malformed eye, with extremely short bodies, or with a permanently extruded pharynx.

**Figure 6.**
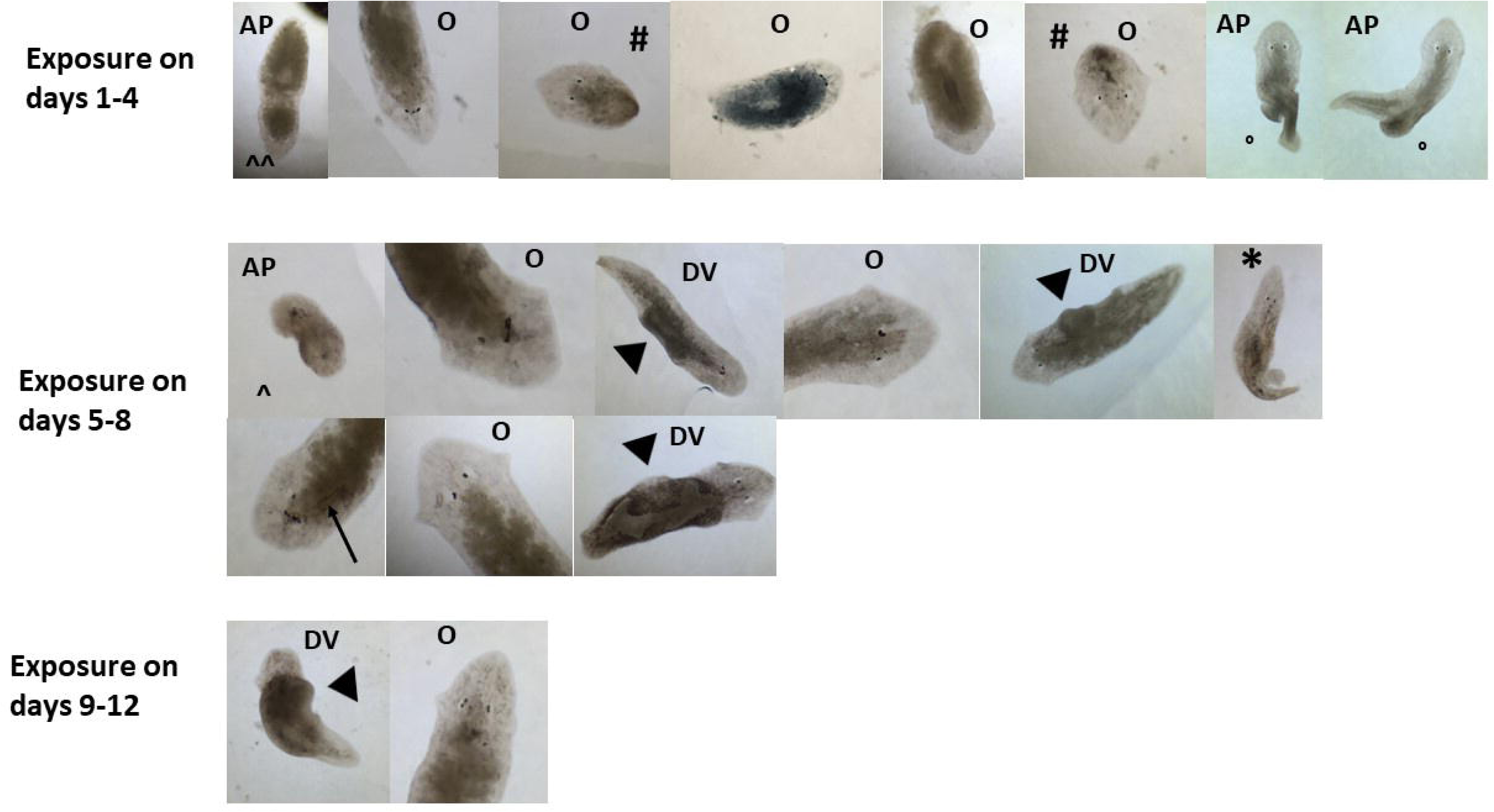
Full range of barium-induced malformations (AP = anteroposterior, DV = dorsoventral, O= other) after exposure on days 1-4, 5-8, and 9-12. Some emerging patterns were noted. For example, dorsal or dorsolateral pigmented “bulges” (arrowheads) were only seen after exposure on days 5-8 and 9-12 (but never on days 1-4), suggesting they are associated with the initiation of BMP expression during the transition from cryptic larva to bilateral embryo. Anteroposterior abnormalities were seen after exposure on days 1-4 and 5-8, including two-headed worms with two eyes (^^) or one eye (^) as well as ectopic anterior bodies embedded on the “main” body of the planaria (°). Interestingly, eye malformations were seen after all exposures. Baby planarias with extremely short bodies were only seen with exposure on days 1-4 (#). Notice the baby planaria with a permanently extruded pharynx (days 5-8, asterisk) and the strange tubular structure emerging from the malformed eye (arrow).

Control experiments with 4 mM NaCl to probe the effects of increased osmolarity/ chloride concentrations showed some toxicity (but no malformations) with continuous exposure (days 0-20). However, 4 mM NaCl was totally innocuous when embryos were exposed during the same stage-specific intervals described in the paper (days 1-4, 5-8, 9-12, 13-16, and 17-20, data not shown).

## Discussion

The concentrations at which barium produces no effects, teratogenic effects, or lethal effects seem to be very close, an effect that could be called “narrow teratogenic index”, and which bears some resemblance to the narrow therapeutic index of certain drugs that work by blocking ion channels, like the anti-epileptic drug phenytoin [20]. This, coupled with “intersubject variability” (differences between individual eggs and embryos, including how quickly they develop) produces some variability in the effects observed (none, lethal or teratogenic) for a specific time window. However, after analysis of multiple samples, a clear pattern emerged: malformations were detected when exposure occurred during the initial cryptic larval stages and during a key morphogenetic event: the transition from radially symmetric cryptic larva to bilaterally symmetric embryo.

Abnormalities in the dorsoventral axis were observed only on the dorsal surface. These were dorsal pigmented “bulges”, seen only after exposure on days 5-8 and 9-12 (but never on days 1-4). Interestingly, these periods coincide with the initiation and upregulation of BMP expression, which starts at approximately day 7 of embryonic development (see figure 4). It has been reported that BMP silencing in adult worms by RNA interference generates less pigmented, dorsal “bulges” in *Dugesia Japonica* and *Schmidtea mediterranea.* Since the ventral side is less pigmented, this was interpreted as an ectopically formed ventral side on top of the worm [21, 22]. In contrast, bioelectric perturbation with barium during days 5-8 and 9-12 of embryonic development induced *more* pigmented dorsal “bulges” in some worms. This suggests that widespread potassium channel inhibition triggers depolarization, hyperactivation of the BMP pathway, and formation of duplicated dorsal tissues on top of the developing worms.

Anteroposterior abnormalities were seen only after barium exposure during days 1-4 and 5-8 of embryonic development (but never on days 9-12) and included two-headed worms (with 2 eyes or 1 eye on each end), and worms with ectopic partial anterior bodies embedded within the “main” body of the planaria. Anteroposterior specification in planaria depends on beta-catenin 1 expression. It has been shown that there is a gradient from the posterior to the anterior end of adult worms, with high beta-catenin 1 specifying posterior identity (23). Interestingly, whole mount in situ hybridization showed that there are two “peaks” of beta-catenin 1 RNA expression during embryonic development: an early one associated with the formation of the cryptic larva (∼days 1-2) and a later one associated with the transition from cryptic larva to bilateral embryo (∼day 7 onwards, figure 4). These two periods coincide with the moments when barium induced anteroposterior abnormalities, suggesting that bioelectric perturbation with barium can modulate these peaks of beta-catenin 1 expression and impact anteroposterior identities. In this regard, voltage reporter dye experiments demonstrated gradients of membrane potential across the anterior-posterior axis of adult planarian flatworms, with the anterior end relatively depolarized and the posterior end relatively hyperpolarized (24) (figure 8). Barium depolarizes cells, suggesting that modification of these voltage gradients during days 1-4 and 5-8 of embryonic development can modulate beta-catenin expression, favoring anterior specification. In this regard, experimental pharmacologic manipulation of these gradients in adult worms has been shown to alter anteroposterior identity. For example, the H,K-ATPase ion pump inhibitor SCH-28080 robustly hyperpolarizes planarian tissues, inhibits head identity in amputated fragments and suppresses ectopic head formation even in the presence of β-catenin RNAi, which induces two-headed worms (24).

In the limited number of samples analyzed, 3 mM barium was lethal or teratogenic during days 1-4 and 9-12 but instead was innocuous or teratogenic during days 5-8, as if a “rheostat” controlling the susceptibility of the embryo to the lethal effects of barium was transiently turned counterclockwise (figure 5). The significance of this observation is unknown, although the period during which it occurs coincides approximately with the transition from radially symmetric cryptic larva to bilaterally symmetric embryo.

In HeLa cells, heterotypic gap junction channels function as voltage-sensitive valves for intercellular signaling [25], and voltage reporter dye experiments demonstrated gradients of membrane potential across the anterior-posterior axis of adult planarian flatworms [11].

These mechanisms seem to play a critical role during planaria embryologic development, as evidenced by the barium-induced malformations of a worm having a second partial body embedded diagonally within the “main” body of the planaria and having a second pair of eyes, or of grossly abnormal two-headed worms with a single eye on each “head” (figure 7B, figure 9), morphological disorganizations that, to the best of the author’s knowledge and despite the impressive plasticity of adult planarias, are not frequently reported in the literature.

**Figure 7.**
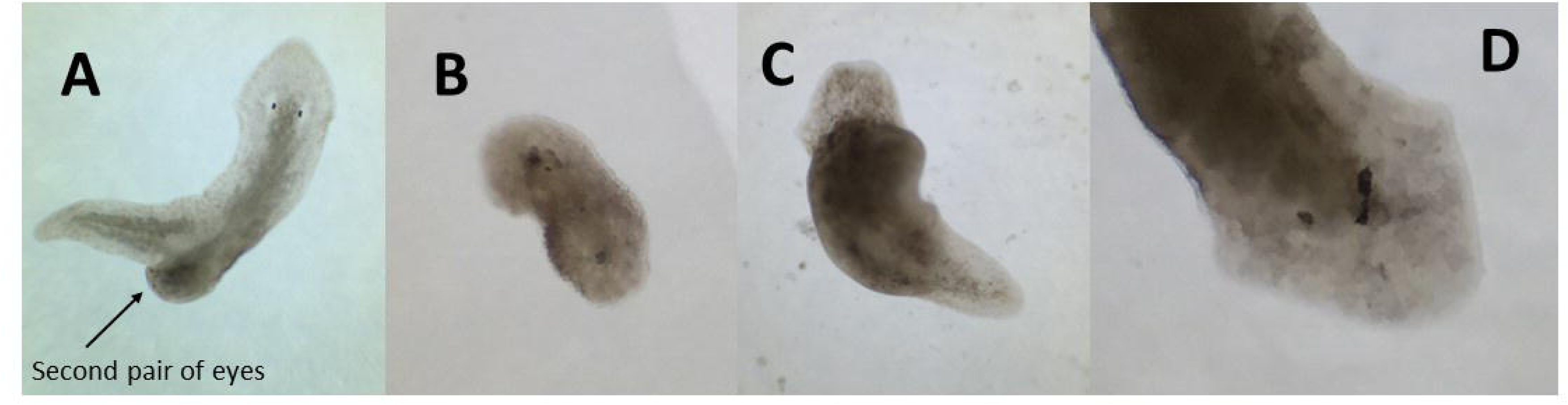
Close-up views of barium-induced malformations in planaria. (A) Malformation after exposure on days 1-4. Notice the second partial body, embedded diagonally within the “main” body of the planaria, with a second pair of eyes. (B) Malformation after exposure on days 5-8: Close-up view of abnormal two-headed worm with a single eye on each “head”. (C) Dorsal pigmented “bulge”. (D) Strange tubular structure associated with malformed eyes.

**Figure 8.**
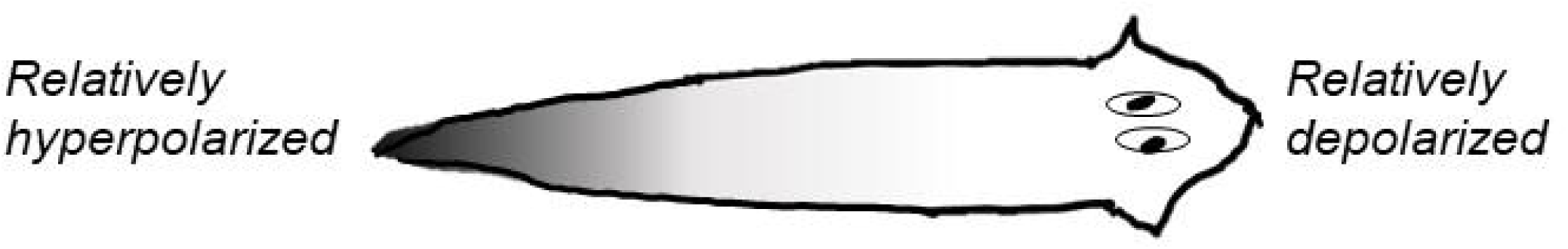
Schematic diagram of voltage gradient in adult planaria (24)

**Figure 9.**
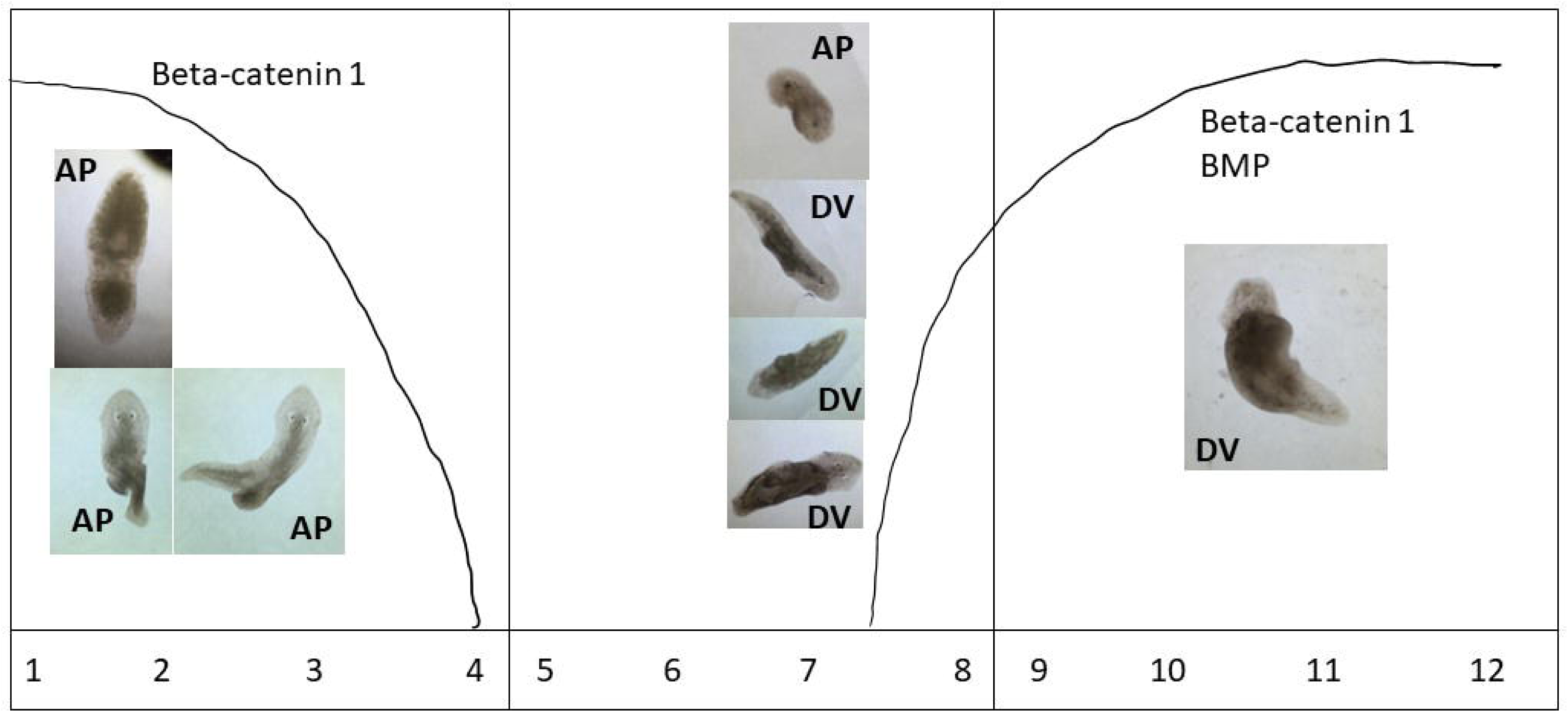
Dorsoventral and anteroposterior abnormalities superimposed on a schematic diagram of relevant gene expression curves. Days are shown on the bottom. Anteroposterior anomalies (anteriorization) occurred mainly after barium exposure on days 1-4, although some were also seen after days 5-8, coinciding with the two beta-catenin 1 gene expression peaks. In contrast, dorsoventral anomalies (pigmented dorsal bulges) were only seen after barium exposure on days 5-8 and 9-12, coinciding with BMP expression.

In other models like Drosophila, inwardly rectifying potassium channels regulate Dpp release during development [26]. Dpp is a member of the BMP family and a key morphogen involved in development. Likewise, Kir2.1 knockout mice show a very similar phenotype to animals in which BMP signaling components have been removed from the cranial neural crest [26].

